# Autophagy degrades immunogenic endogenous retroelements induced by 5-azacytidine in acute myeloid leukemia

**DOI:** 10.1101/2022.12.02.518683

**Authors:** Nandita Noronha, Chantal Durette, Bianca E Silva, Justine Courtois, Juliette Humeau, Allan Sauvat, Marie-Pierre Hardy, Krystel Vincent, Jean-Philippe Laverdure, Joël Lanoix, Frédéric Baron, Pierre Thibault, Claude Perreault, Gregory Ehx

## Abstract

The hypomethylating agent 5-azacytidine (AZA) is the first-line therapy for acute myeloid leukemia (AML) patients unfit for intensive chemotherapy. Evidence suggests that the anti-tumor effect of AZA results partly from T-cell cytotoxic responses against MHC-I-associated peptides (MAPs) whose expression is induced by hypomethylation. Through a proteogenomic approach, we analyzed the impact of AZA on the transcriptome and MAP repertoire of four AML cell lines and validated salient findings in the transcriptome of 437 primary AML samples. We demonstrate that AZA caused pleiotropic changes in AML cells via perturbation of transcription, translation, and protein degradation. Overall, 1,364 MAPs were upregulated in AZA-treated cells, including several cancer-testis antigens. Increased MAP abundance was due to the upregulation of corresponding transcripts in a minority of cases and post-translational events in most cases. Furthermore, AZA-induced hypomethylation increased the abundance of numerous transcripts, of which 38% were endogenous retroelements (EREs). Upregulated ERE transcripts triggered innate immune responses but were degraded by autophagy and not processed into MAPs. Autophagy resulted from the formation of protein aggregates caused by AZA-dependent inhibition of DNMT2, a tRNA-methyl transferase enzyme. We found that autophagy inhibition had a synergistic effect with AZA on AML cell proliferation and survival, increased ERE levels and triggered pro-inflammatory responses. Finally, autophagy gene signatures were associated with a lower abundance of CD8^+^ T-cell markers in AML patients expressing high levels of EREs. Altogether, this work demonstrates that the impact of AZA is regulated at several levels and suggests that inhibiting autophagy could improve the immune recognition of AML blasts in patients.

## 1. INTRODUCTION

Acute myeloid leukemia (AML) is the most common acute leukemia in adults, with an overall 5-year survival below 30%. Standard therapy involves intensive chemotherapy with a ‘7+3’ regimen of cytarabine and anthracycline. Although AML is a heterogeneous disease, aberrant genomic methylation (hypermethylation in particular (1,2)) is a hallmark of AML blasts. Therefore, hypomethylating agents (HMAs) such as 5-azacytidine (azacitidine, AZA) and 5-aza-2’-deoxycytidine (decitabine, DAC) are used as first-line therapy for AML patients unfit for intensive chemotherapy (3). AZA is also used in maintenance therapy for fit patients without an *FLT3* mutation (3). However, only 18–47% of patients respond to these therapies, stressing the need to improve therapy efficacy, possibly by combining them with other pharmacologic agents (4,5).

AZA and DAC are cytidine nucleoside analogs that incorporate into genomic DNA during replication in the prophase of mitosis (6). High concentrations of AZA and DAC exert cytotoxic effects by inducing DNA double-strand breaks. However, at low concentrations, they act as suicide substrates for DNA methyltransferases (DNMTs) 1 and 3, leading to their degradation and the DNA demethylation of daughter cells. AZA differs from DAC by its ability to incorporate into RNA and DNA, thus inhibiting DNMT2, a transfer RNA (tRNA) methyltransferase (7). While both HMAs have similar response rates in AML (8), only AZA significantly improves overall survival compared with conventional care regimens in phase III randomized trials (9,10). Therefore, AZA is currently FDA-approved as a first-line treatment in AML (11).

In addition to their cytotoxic and demethylating effects, HMAs may mediate anti-leukemic activities by promoting effector immune cells to recognize malignant blasts. Specifically, HMAs enhance the expression of transcripts coding for cancer-testis antigens (CTAs) (12,13). CTA genes are normally silenced by genomic methylation and code for antigens deemed immunogenic because they are not expressed in normal MHC-positive somatic cells (14). Accordingly, some studies evidenced that HMAs promote cytotoxic T-cell (CTL) activity (15), while others demonstrate a specific cytotoxic activity against CTAs (13,16,17). These studies suggest that HMA-induced CTAs can promote anti-leukemic CD8 T-cell reactions by generating immunogenic MHC-I-associated peptides (MAPs).

Along with CTAs, HMAs promote the expression of endogenous retroelements (EREs) (18,19). EREs are highly repetitive sequences that are remnants of transposable elements incorporated into the human genome millions of years ago (20). EREs represent ~45% of the human genome and can be separated into LINEs and SINEs (long and short interspersed elements, respectively) and LTRs (long terminal repeats), the latter of which includes endogenous retroviruses (ERVs). EREs are epigenetically silenced mainly by genomic methylation in normal somatic cells (21) and dysregulated ERE expression is associated with several pathologic conditions, including autoimmunity, inflammatory disorders, aging, and cancer (22). HMA-induced ERE overexpression in solid cancers leads to viral mimicry and concomitant innate immune response (23,24). Moreover, we and others have demonstrated that in addition to being expressed, EREs are capable of being presented by MHC-I molecules and serve as immunogenic tumor antigens, notably in AML (25–28). While EREs are perfect candidates for generating immunogenic MAPs following HMA treatment, there is a lack of robust evidence to support that HMAs enhance their MAP presentation (and subsequent CTL responses) in AML.

T cells recognize MAPs, not transcripts. Hence, available transcriptomic studies do not allow inferences on the MAP repertoire of AZA-treated cells. Herein, we sought to directly evaluate AZA’s capacity to enhance the presentation of CTAs and ERE-derived MAPs in AML. We discovered that AZA promotes the expression of multiple CTA-derived, but not ERE-derived MAPs. Mechanistically, we propose that the lack of ERE-derived MAPs is due to AZA-induced autophagy of ERE transcripts.

## 2. RESULTS

### Low-dose AZA leads to delayed, transient ERE expression in AML cell lines

To investigate the effects of AZA on the immunopeptidome of AML, we selected four AML cell lines (THP-1, MOLM-13, SKM-1, and OCI-AML3) belonging to aggressive FAB types (M4/M5) and together covering different but frequent mutational statuses (MLL-AF9, FLT3-ITD, TET2 (L1418fs), and NPM1c+DNMT3A (R882C), respectively). As we wished for our immunopeptidomic analyses to reveal the effects of AZA independently of cytotoxic activity, we established a protocol that allowed DNMT1 degradation and genomic DNA demethylation without affecting viability or inducing DNA damage responses. The most desirable AZA doses were 0.25 μM (for MOLM-13 and SKM-1) and 0.5 μM (for THP-1 and OCI-AML3) as they sufficiently reduced cell growth, genome methylation, and DNMT1 expression without inducing cytotoxic effects (Figure 1A, Supplementary Fig. S1).

**Figure 1.**
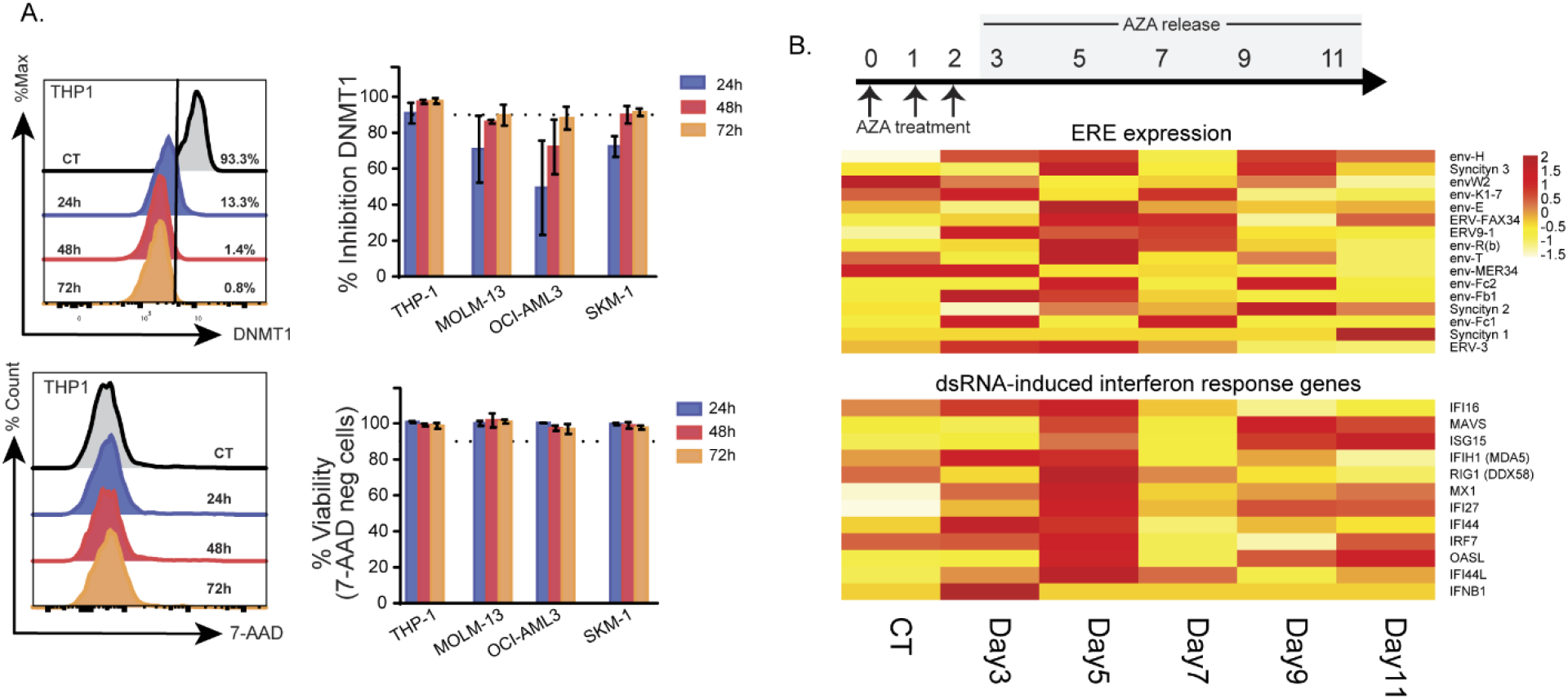
Low-dose AZA treatment leads to delayed, transient ERE and dsRNA-induced interferon gene expression in AML cell lines. **A**, Low AZA doses were added to four AML cell lines daily for three days (0.25 μM: MOLM-13 and SKM-1; 0.5 μM: THP-1 and OCI-AML3) and DNMT1 inhibition (upper panel) and cell viability were monitored by flow cytometry using 7-AAD (lower panel). Dotted lines represent 90% DNMT1 inhibition/viability in the upper and lower panel respectively. The left panels depict representative histograms of THP-1 cells, while the right panels depict bar plots summarizing the percentage of expression/staining (as indicated in figures) of all four AML cells. Percentages were calculated by comparing AZA-treated cells to the control cells. **B**, Low AZA doses (0.5 μM) were added to THP-1 cells daily for three days, after which AZA treatment was released by washing and expanding cells in the absence of AZA. Cells were collected for RNA-seq at the time points highlighted in grey. Heatmap comparing RNA expression levels of EREs induced by AZA (upper panel) and genes involved in dsRNA-induced viral response (lower panel) identified in previous studies (23,24) of control (day 3) and AZA-released cells at the time points indicated.

Since previous studies in ovarian and colorectal cancer demonstrated that AZA leads to delayed ERE expression (23,24), we sought to identify the time point post-AZA discontinuation with the highest ERE expression. Using THP-1 as a model, we performed RNA sequencing (RNA-seq) every 48h from day 3 to 11 post-AZA discontinuation. We then quantified the expression of multiple ERE transcripts upregulated by AZA (24). We also assessed the expression of genes involved in dsRNA-induced interferon signaling in response to AZA-induced ERE expression (23) and observed that, together with ERE transcripts, they reached maximum expression around day 5 (72h after the last AZA treatment; Figure 1B). qPCR analyses further validated that the effect of AZA on ERE and dsRNA-induced immune response genes was maximal on day 4 (48h after the last AZA treatment; Supplementary Fig. S2). Thus, we treated the four AML cell lines with these optimal AZA doses, administered three times at 24h intervals (0h, 24h, and 48h), and harvested the cells on day 4 to perform RNA-seq and mass spectrometry (MS) analyses.

### Proteogenomic characterization of the immunopeptidome

We have previously demonstrated that a significant fraction of the AML immunopeptidome derives from non-exonic genomic regions such as introns, EREs, or intergenic regions (28). Typically, MS search engines rely on reference protein databases (such as Uniprot) to match each acquired MS/MS spectrum to a peptide sequence. However, these databases do not contain non-exonic sequences, and building a personalized database containing all genomic sequences would generate an unmanageable database for search engines. Therefore, we used a proteogenomic approach to create MS databases containing only MAP sequences corresponding to the RNA transcripts expressed by our cell lines. These customized cell-line-specific MS databases had manageable sizes for the PEAKS search engine (Figure 2A and Supplementary Fig. S3).

**Figure 2.**
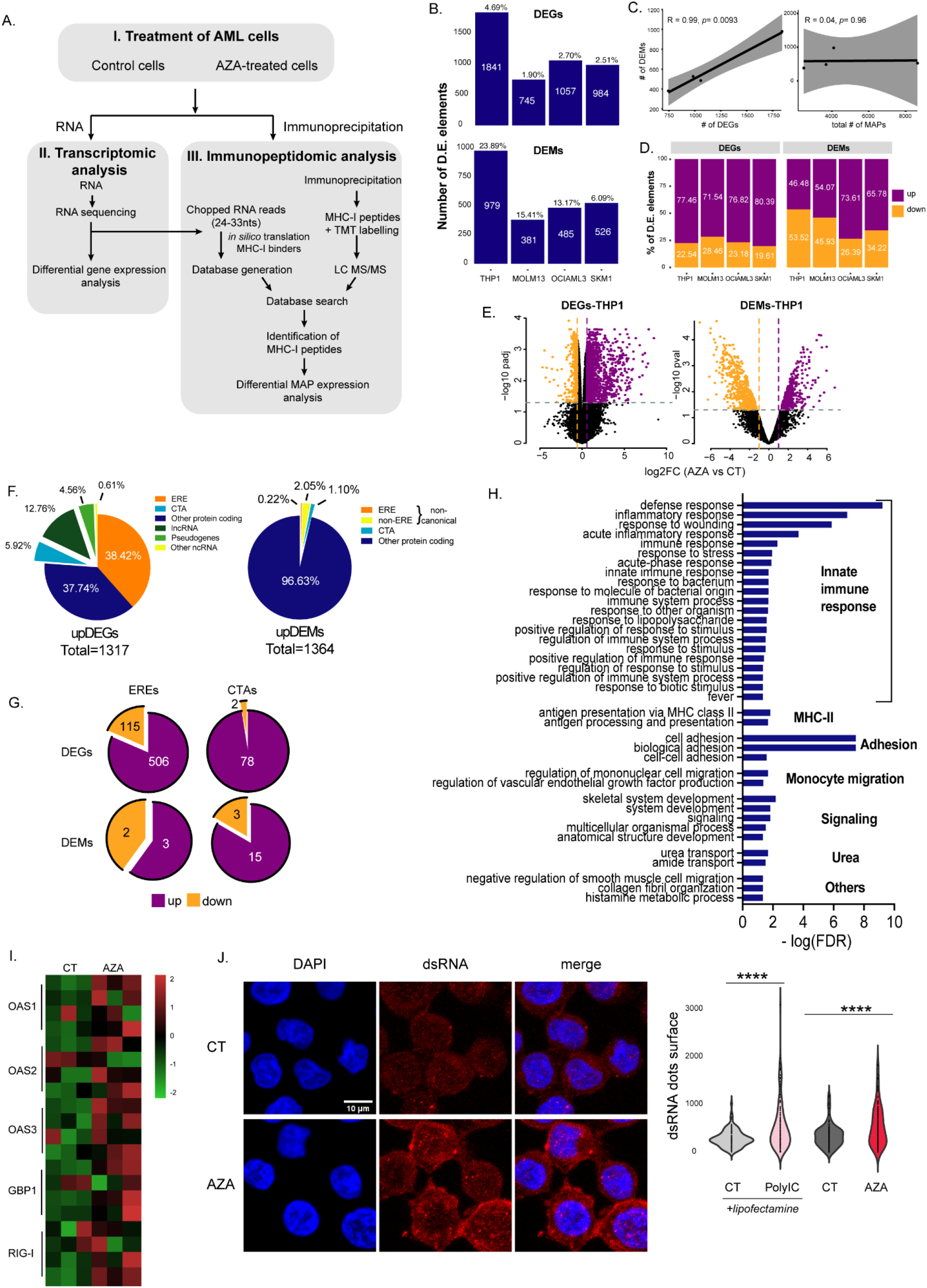
Proteogenomic characterization of AZA-mediated changes shows upregulation of MAPs derived from CTA but not from EREs. **A**, Schematic representation of the study design for RNA-seq and MS analyses. **B**, The total number of differentially expressed (D.E.) genes and MAPs (DEGs and DEMs, respectively) varies across cell lines. The numbers above the bars indicate the percentage of total genes or MAPs that were DEGs or DEMs, respectively. **C**, Pearson correlation between the number of DEMs and DEGs (left panel) or DEMs and MAPs (right panel). Each dot corresponds to a cell line. **D**, Percentage of D.E. elements up- or downregulated across cell lines. **E**, Representative volcano plots of DEGs and DEMs between AZA (violet) and untreated (gold) THP-1 cells. **F**, Pie charts depicting the percentage of biotypes of upregulated transcripts (left) and MAPs (right). The total number of upregulated D.E. elements is indicated below the pie charts. **G**, Pie charts depicting the number of up- and downregulated EREs (left) and CTAs (right) belonging to DEGs (upper panel) or DEMs (lower panel) fractions. **H**, Histogram depicting GO-term analysis of the most significantly enriched biological processes associated with upregulated DEGs across all cell lines after AZA treatment. **I**, Heatmap of genes involved in anti-dsRNA responses. **J**, Representative images (left) and quantification (right) of dsRNA signals in THP-1 cells from microscopy images (two independent experiments). THP-1 cells transfected with 10 μg/ml polyinosinic:polycytidylic acid using lipofectamine (PolyIC+lipofectamine) were used as a positive control and were compared with cells treated with lipofectamine alone (CT+lipofectamine); (unpaired t-test; **** p < 0.0001).

Precisely, the RNA-seq reads were first chopped into shorter sequences (k-mers, of 24–33 nucleotides) corresponding to the different possible MAP lengths (8–11 residues). These k-mers were then filtered based on inter-sample sharing and abundance since abundant transcripts have higher chances of generating MAPs (28,29). K-mers possibly deriving from sequencing artifacts or lowly abundant polymorphisms were removed. The resulting k-mers were then *in silico* translated into peptide sequences, and those present in the canonical proteome were discarded. Finally, peptides were evaluated for their predicted capacity to bind MHC-I allotypes of their respective cell line, and binders were concatenated with the canonical proteome to generate MS databases ranging between 250 and 350 Mb (sizes tolerated by MS search engines (28)).

In parallel to MS, we performed a differential expression analysis of annotated proteincoding genes and ~4.2×10^6^ ERE regions reported in the Repeatmasker annotations (30). Differential abundance analyses were also performed on MS data to correlate transcript expression and MAP presentation. These analyses revealed that the number of elements differentially expressed by AZA-treated cells varied across cell lines, with THP-1 being the most sensitive (Figure 2B, Supplementary Table S1). Overall, the proportion of differentially expressed MAPs (DEMs) was five times greater than that of differentially expressed transcripts (DEGs): 6–23% vs. 1.9–4.69%, respectively. Notably, the number of DEMs per cell line correlated almost perfectly with the number of DEGs rather than the total number of MAPs per cell line, suggesting that transcriptomic alterations were reflected in the immunopeptidome (Figure 2C). However, the directionality of differential expression differed for MAPs and transcripts. While most DEGs (>70%) were upregulated by AZA, this was not the case for DEMs (Figures 2D-E, Supplementary Table S2). This means that as with other drugs (31), changes in the immunopeptidome post-AZA treatment result from differential mRNA expression and post-translational events.

### AZA-induced EREs do not generate MAPs but trigger innate immune responses

Next, we focused on variations in CTA and ERE expressions at the mRNA and MAP levels. As expected, we observed a striking upregulation of ERE transcripts, representing ~38% of upregulated DEGs (Figure 2 F-G, Supplementary Table S3). In contrast, EREs represented only 0.22% of upregulated DEMs, meaning that AZA-induced ERE transcripts were not processed adequately for MHC-I presentation. Due to the delay between AZA treatment and ERE induction (Figure 1B), we repeated our immunopeptidomic analyses at a later time point (day 7) on THP-1 (the cell line with the highest AZA-induced EREs) and did not observe higher ERE MAP presentation (Supplementary Fig. S4). In contrast with EREs, we observed lower discrepancies in proportions of CTA-derived elements among upregulated DEGs and DEMs, and they were still observed on day 7. Among the upregulated DEMs, 152 (~10%) were AZA-specific (i.e., presented by all three AZA-treated replicates but undetected in control cells; Supplementary Fig. S5A). MAPs induced *de novo* by AZA contained CTAs but not ERE MAPs (Supplementary Fig. S5B). We conclude that at the immunopeptidomic level, AZA upregulates the expression of CTA MAPs (some of which are AZA-specific) but not ERE MAPs. The latter point was striking, considering the dramatic upregulation of ERE DEGs (38% of all DEGs) following AZA treatment.

Because EREs are remnants of ancient viruses, they can induce innate immune responses triggered by double-stranded RNA (dsRNA). Accordingly, we investigated whether AZA-induced EREs would trigger such responses. Gene ontology (GO) analysis performed on upregulated DEGs revealed that multiple innate immune responses and inflammatory responses were triggered in AZA-treated cells (Figure 2H). Specifically, we observed that OAS1, OAS2, OAS3, GBP1, and RIG-I – five genes instrumental in anti-dsRNA responses – were expressed at higher levels in AZA-treated cells than in controls (Figure 2I). Accordingly, using microscopy, we observed greater amounts of dsRNA in AZA-treated cells than in controls (Figure 2J). These data suggest that the dramatic upregulation of ERE transcripts induced by AZA leads to dsRNA formation, thereby triggering innate anti-viral immune responses.

### AZA-induced EREs correlate with innate immune responses in primary AML

In cancer cells, innate immune responses benefit the host because they can initiate cancer cell apoptosis and increase their adjuvanticity (32). We, therefore, asked two questions: i) what is the profile of EREs induced by AZA, and ii) can this profile be detected in primary AML samples? We first noted that ERE induction by AZA in AML cell lines was not random. Using ERE distribution in the genome as a reference, we observed that AZA selectively upregulated two classes of EREs: LINEs and LTRs (Figure 3A). The fact that repression of SINEs depends mainly on histone methylation rather than DNA methylation (33) can explain why AZA did not induce SINE expression.

**Figure 3.**
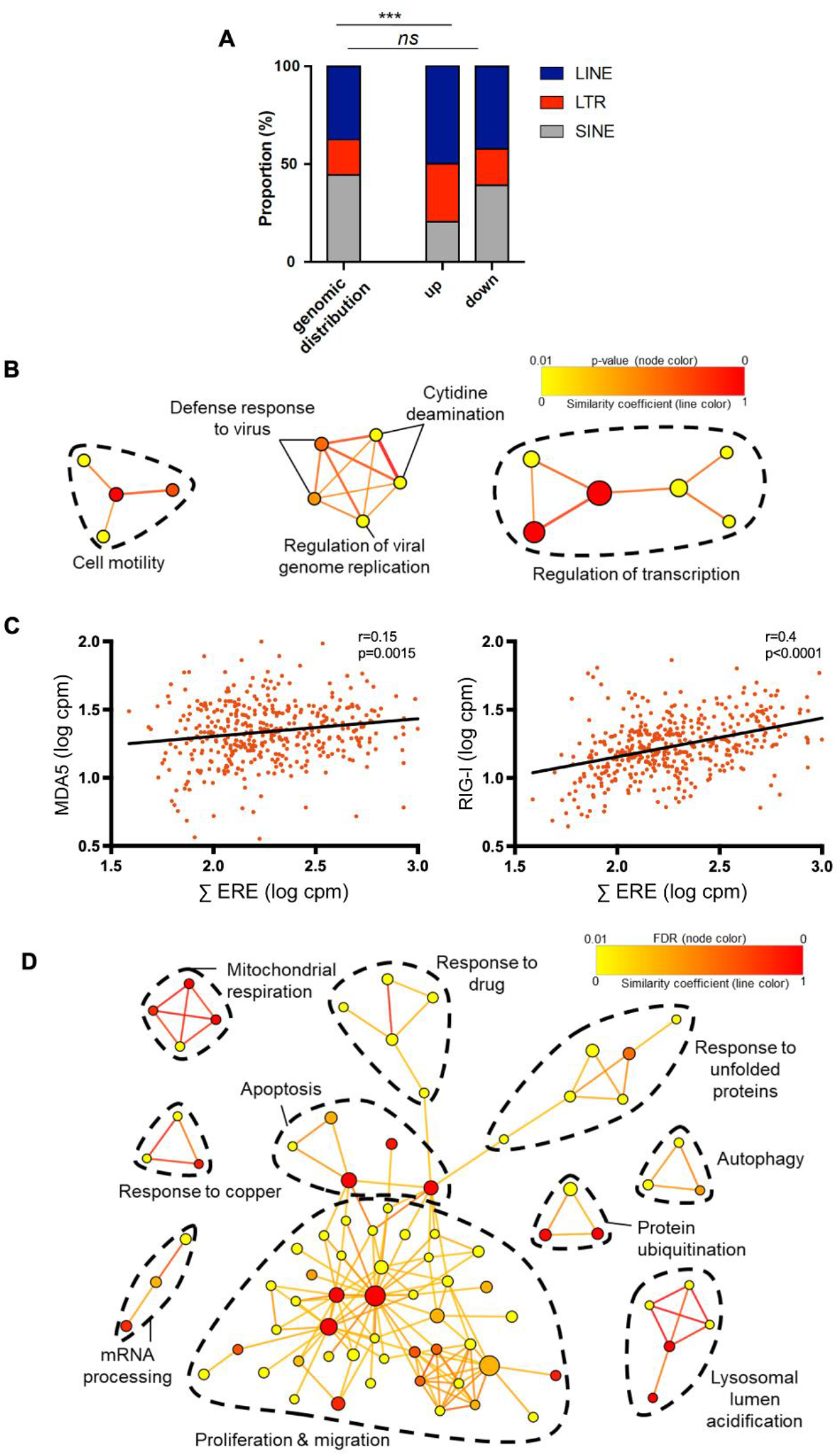
AZA-induced EREs trigger innate immune responses. **A**, Stacked bar plots of ERE group distribution at the genomic and transcriptomic levels for up- and downregulated AZA-altered ERE sequences. **B**, Network analysis of GO-terms in AML patients (Leucegene cohort; n=437) expressing high levels of EREs induced by AZA in our four cell lines. **C**, Pearson correlation between genes involved in anti-dsRNA responses (MDA-5 and RIG-I) and AZA-induced EREs in AML patients. **D**, Network analysis of GO-terms enriched in patients expressing low levels of AZA-induced EREs. In B, and D, the line color reflects the similarity coefficient between connected nodes. Node color reflects the false discovery rate (FDR) of the enrichment. Node size is proportional to gene set size.

Using the previously published RNA-seq data of the Leucegene cohort (primary AML samples from 437 patients), we quantified the expression of the 506 ERE transcripts significantly upregulated by AZA in our AML cell lines. Patients were segregated based on their cumulative expression of these EREs, and patients expressing above-median levels were compared with those expressing below-median levels. DEGs and GO analyses revealed that high levels of AZA-induced EREs were associated with upregulated defense responses against viruses (Figure 3B). Furthermore, ERE expression levels significantly correlated with the expression of two critical dsRNA response regulators: RIG-I and MDA5 (Figure 3C), supporting the notion that AZA-induced EREs trigger innate immune responses *in vitro* and *in vivo*.

To complement our previous analysis, we performed GO analyses on genes downregulated by AML patients expressing high levels of AZA-induced EREs. This showed that multiple pathways controlling proliferation were downregulated, suggesting that high ERE expression (and anti-dsRNA response) could impact the growth of AML blasts (Figure 3D). Unexpectedly, we also observed that many GO terms related to protein degradation/catabolism and autophagy were significantly downregulated in these patients. Since ERE RNAs can trigger autophagy (34) and be degraded by the autophagic process (35), we hypothesized that enhanced autophagy in low-ERE expressing blasts could protect them from the deleterious effects that EREs have on their proliferation. This idea was explored in-depth in our next series of experiments.

### AZA molds the immunopeptidome and induces protein aggregation through DNMT2 inhibition

We next sought to explore AZA-induced changes in the immunopeptidome for two reasons: First, because of the disconnect between the number of ERE DEGs and ERE DEMs (Figure 2F) and second, because our initial MAPs of interest (CTAs and EREs) represented only a fraction of AZA-induced MAPs (Figure 2F). To this end, we assessed the global impact of transcriptomic variations on the immunopeptidome using BamQuery, a computational tool that quantifies the RNA expression of any MAP of interest, including those derived from non-annotated genomic regions (36). Most AZA-altered DEMs displayed no change at the RNA level (Figure 4A). Nevertheless, among DEMs coded by DEGs, RNA upregulation strongly correlated with convergent upregulation of the corresponding DEMs (Figure 4A). This was not the case for downregulated transcripts. Moreover, fold changes in RNAs generating upregulated DEMs were significantly higher than those in downregulated DEMs (Figure 4B). We conclude that transcript upregulation has a modest but genuine impact on the abundance of corresponding DEMs.

**Figure 4.**
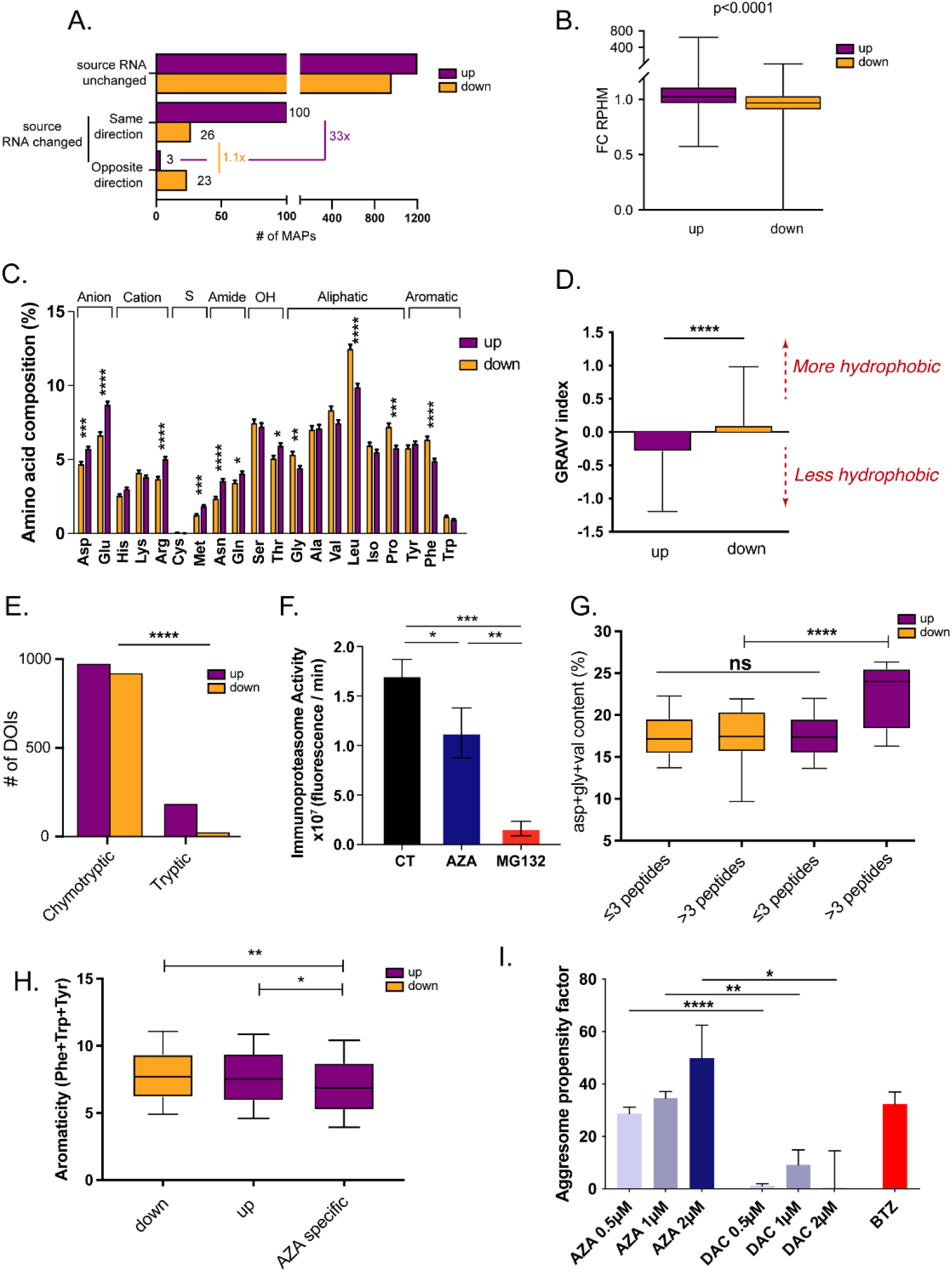
AZA molds the immunopeptidome through DNMT2 inhibition. **A**, Bar plots depicting the number of up- and downregulated DEMs with no change in source RNA expression or being altered in the same or opposite direction as their source RNA. **B**, Fold changes in RPHM expression of source transcripts generating up- and downregulated DEMs. **C**, Amino-acid composition of up- and downregulated DOIs. **D**, Hydrophobicity of up- and downregulated DOIs assessed by their GRAVY index. Scores >0 reflect higher hydrophobicity. **E**, Number of DOIs associated with chymotryptic or tryptic activities based on their C-termini amino acid composition (Fisher’s exact test). **F**, Immunoproteasome activity in OCI-AML3 cells after AZA treatment. MG132, a proteasome inhibitor, was used as a negative control (unpaired t-test). **G**, Proportion of DNMT2-target amino acids (glycine, valine, and aspartic acid) in proteins having generated MAPs among the up- or downregulated DEM fractions. **H**, Aromaticity (frequency of Phe, Trp, and Tyr residues) of proteins having generated AZA-specific (identified only in AZA condition) MAPs or up- and downregulated DEMs. **I**, Quantification of protein aggregates induced with increasing concentrations of AZA and DAC in OCI-AML3 cells. Bortezomib (BTZ) was used as a positive control (unpaired t-test; **** p< 0.0001, *** p< 0.001, ** p< 0.01, * p< 0.05).

We next focused on DEMs whose source RNA fold-change did not explain their immunopeptidomic fold-change (DEMs of Interest, DOIs). To gain insights into co- or post-translational events conducing to immunopeptidomic alterations in AZA-treated cells, we started by analyzing the residue composition of DOIs. We found that upregulated DOIs contained more polar residues than downregulated DOIs (Figure 4C). Accordingly, these MAPs presented a lower overall hydrophobicity (Figure 4D). Hydrophobic residues are the preferential cleavage sites of proteasomes, particularly immunoproteasomes (37). MAP generation by constitutive proteasomes depends mainly on their tryptic and chymotryptic-like activities, and chymotryptic-like activity is further amplified in immunoproteasomes (38,39). Hence, we assessed how protease activity contributes to the immunopeptidome by examining the C-terminal residue of each DOI and observed that DOIs derived more frequently from tryptic cleavage (Figure 4E). Accordingly, AZA treatment significantly reduced immunoproteasome activity (Figure 4F, Supplementary Fig. 6A).

Typically, alterations in proteasomal activity are associated with disrupted protein homeostasis (40,41). To investigate whether alterations in protein homeostasis were responsible for perturbed proteasomal activity in AZA-treated cells, we examined the residue composition of proteins that generated DOIs. Assuming that proteins generating multiple DOIs were degraded more actively than those generating a single DOI, we correlated the number of generated DOIs for each protein with the frequency of each residue in the considered protein. This analysis showed that aspartic acid (Asp) and glycine (Gly) had the strongest positive correlation with the number of upregulated DOIs (Supplementary Table S4). Interestingly, the tRNAs of Asp and Gly are stabilized by DNMT2, a tRNA-methyl transferase enzyme inhibited by AZA (42,43). A targeted analysis comparing the frequency of Asp, Gly, and valine (Val, the third amino acid whose tRNA is methylated by DNMT2) revealed that proteins generating more than three upregulated DOIs presented significantly higher cumulative frequencies of Asp, Gly, and Val than those generating less than three upregulated DOIs (Figure 4G). Demethylated tRNAs are susceptible to ribonuclease cleavage and fragmentation (44). Therefore, we hypothesized that AZA-mediated DNMT2 inhibition results in an insufficiency of Asp, Gly, and Val tRNAs and leads to a decrease in protein synthesis, ribosomal stalling, and consequent protein aggregate generation during translation of proteins rich in the aforementioned residues. Accordingly, AZA-specific peptides (technically the most upregulated DOIs) were generated from proteins with a significantly lower proportion of aromatic residues, a feature often associated with less efficient protein folding (Figure 4H) (45). Finally, we experimentally quantified protein aggregates in AZA-treated cells and discovered that AZA induced protein aggregate accumulation in a dose-dependent manner (Figure 4I, Supplementary Fig. 6B). Importantly, DAC, a hypomethylating drug that does not inhibit DNMT2, did not induce the formation of protein aggregates.

### Autophagy degrades AZA-induced EREs

Given the inverse correlation between autophagy-related GO terms and EREs in AML patients (Figure 3B) and the generation of protein aggregates by AZA (Figure 4I), we evaluated whether AZA induced autophagy. Twenty-four hours of treatment with AZA resulted in a dose-dependent induction of autophagy (Figure 5A). Interestingly, this was not observed with DAC, suggesting that autophagy induction is dependent on protein aggregates generation resulting from DNMT2 inhibition. As EREs were previously suggested to trigger autophagy (34), we evaluated whether DAC induced the same EREs as AZA. Using data reported by Pappalardi *et al*. (46), we observed that DAC expressed AZA-induced EREs in a dose-dependent manner (Figure 5B). This suggests that the autophagy induced in AZA-treated (but not in DAC-treated) cells results from DNMT2 inhibition rather than ERE induction. Nevertheless, we observed an inverse correlation between EREs and two well-established autophagy markers, ATG3 and SQSTM1, which was already observed in AML patients at diagnosis (Figure 5C). The latter finding suggests that autophagy does not need to be induced by AZA to degrade EREs.

**Figure 5.**
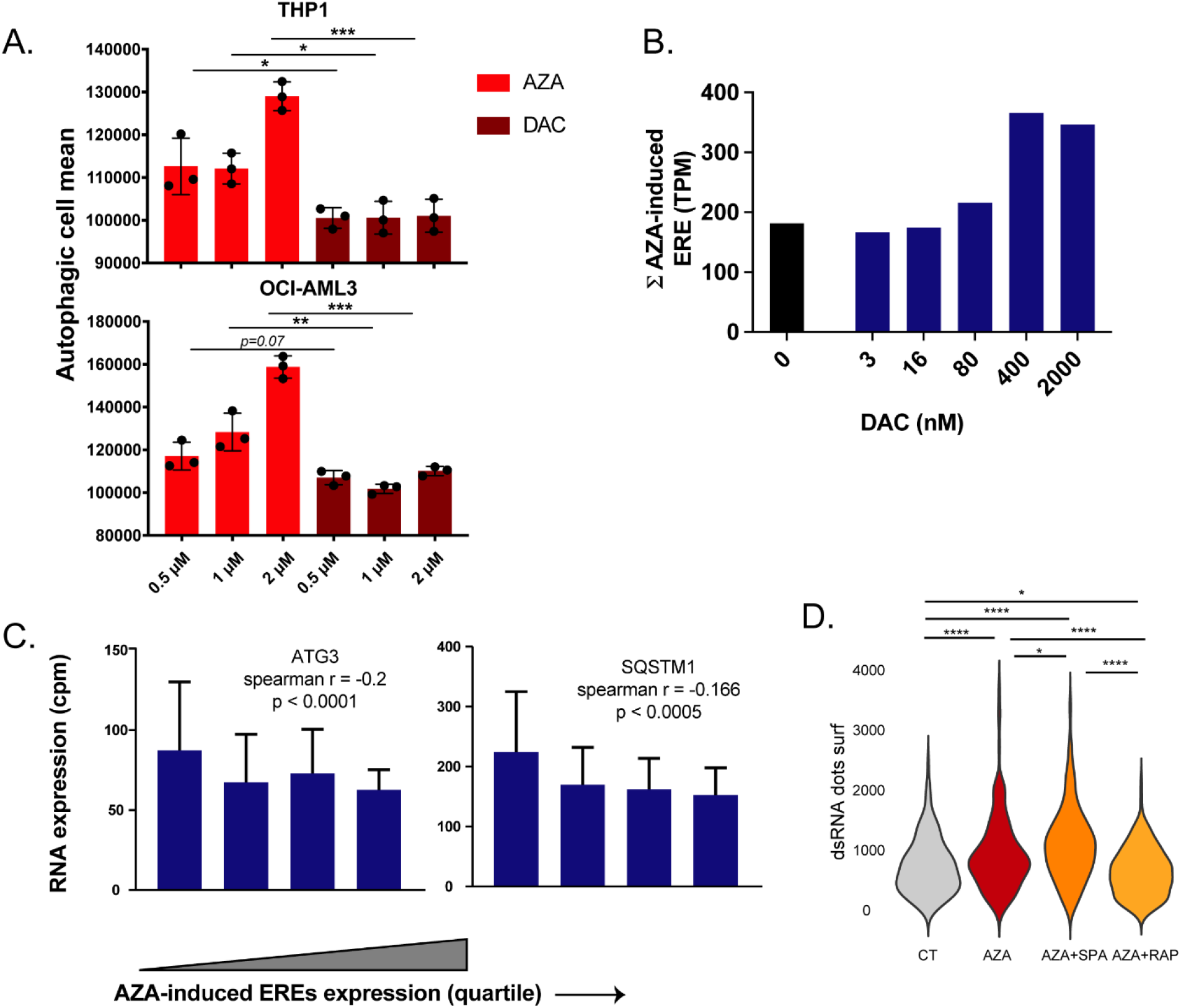
Autophagy degrades AZA-induced EREs. **A**, THP-1 and OCI-AML3 cells undergoing autophagy were assessed with increasing doses of AZA and DAC by flow cytometry using specific autophagy detection fluorescent probes. **B**, EREs induced by AZA in our cell lines were quantified in published RNA-seq data of DAC-treated THP-1 cells (47). **C**, Bar plots of mean RNA expression (in cpm) of key autophagy genes (ATG3 and SQSTM1) in Leucegene AML patients segregated into quartiles based on AZA-induced ERE expression. Spearman correlations were computed without this segregation. **D**, Quantification of dsRNA signals measured from microscopy images. CT: 0.1% DMSO treated with lipofectamine; AZA+SPA: AZA and spautin-1; AZA+RAP: AZA and rapamycin (unpaired t-test; * p< 0.05, **** p< 0.0001).

Next, we sought to verify this autophagy-dependent degradation of EREs. We treated THP-1 cells for 72h with AZA alone or AZA combined with an autophagy inducer (rapamycin) or inhibitor (spautin-1). Ninety-six hours after the initiation of the treatment, the levels of dsRNAs were examined by fluorescence microscopy. As shown in figure 5D, autophagy inhibition significantly increased the levels of dsRNAs compared to AZA alone, while autophagy activation decreased them. Altogether, these results demonstrate that autophagy contributes to the degradation of EREs.

### Autophagy inhibition synergizes with AZA and could increase AML immunogenicity

Finally, we examined whether inhibiting autophagy would augment the anti-AML effect of AZA. THP-1 and OCI-AML3 cells were treated for 72h with AZA and/or spautin-1 (which inhibits autophagy and suppresses the unfolded protein response (48)), and their survival and cell counts were evaluated 24h after discontinuing treatment. A synergistic effect between AZA and spautin-1 was observed for proliferation and cell death (Figure 6A, Supplementary Fig.7A). While spautin-1 alone reduced proliferation, it did not kill the cells (Supplementary Fig. 7B, C), suggesting that autophagy acts as a survival mechanism upon AZA treatment.

**Figure 6.**
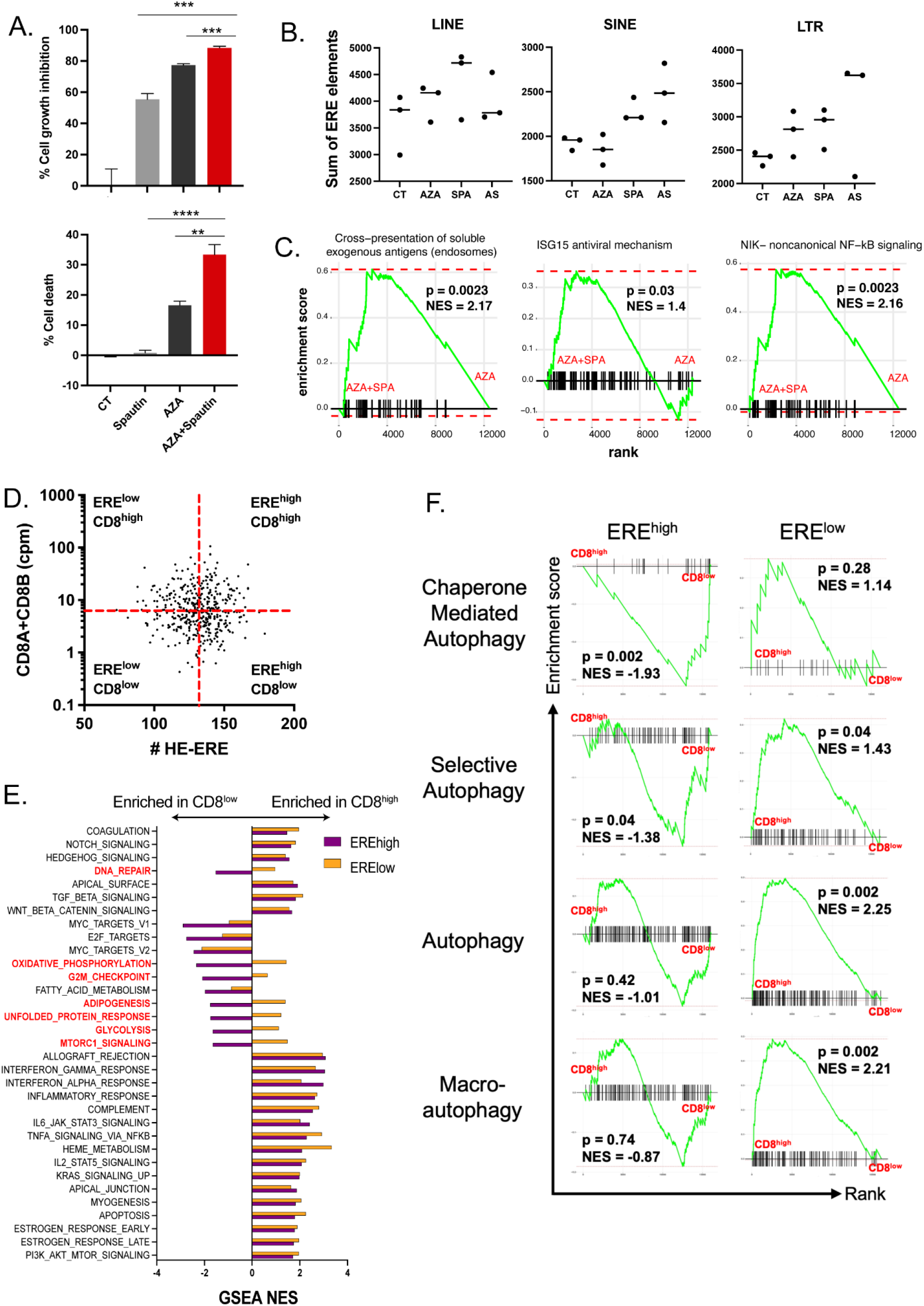
Autophagy inhibition synergizes with AZA and could increase AML immunogenicity. **A**, Cell growth inhibition and cell death (7-AAD) of THP-1 cells treated either with AZA (1 μM), spautin-1 (5 μM), or both. Control cells were treated with 0.1% DMSO; two independent experiments (unpaired t-test; **** p< 0.0001, *** p< 0.001, ** p< 0.01). **B**, Sum of ERE transcripts (in cpm) separated into LINE, SINE, and LTR families in THP-1 cells, treated as indicated. CT: 0.1% DMSO; SPA: spautin-1; AS: AZA and spautin-1. **C**, GSEA comparison of indicated gene sets between THP-1 cells treated with AZA or AZA + spautin-1. NES, normalized enrichment score. **D**, Scatterplots of Leucegene AML patients based on their RNA expression of CD8A+CD8B (cpm) vs. their count of highly expressed AZA-induced EREs (HE-EREs: # of AZA-induced EREs whose expression is above their median expression across all patients). **E**, Bar plots indicating normalized enrichment scores for hallmark gene sets between CD8^high^ vs. CD8^low^ Leucegene patients within ERE^high^ and ERE^low^ groups (defined in D panel). **F**, GSEA of the indicated REACTOME gene sets in the indicated comparisons. NES, normalized enrichment score.

Next, we investigated the molecular effects of autophagy inhibition combined with AZA. THP-1 cells treated with low AZA doses and/or IC50 concentrations of spautin-1 were analyzed by RNA-seq, and as expected, we observed that spautin-1 alone increased the levels of all three ERE families (Figure 6B). Furthermore, ERE levels tended to be higher (except for LINEs) when spautin-1 was combined with AZA. In contrast, AZA-induced EREs in these cells were at the same levels as in control cells (Supplementary Fig. 8B), possibly due to the drastic inhibition of proliferation (Supplementary Fig. 8A) mediated by the combined treatment (AZA’s DNA demethylating activity relies on active cell division for AZA to integrate into genomic DNA). Nevertheless, a pathway enrichment analysis on genes significantly upregulated in AZA+Spautin-1 vs. AZA-treated cells revealed the enrichment of multiple pathways related to antigen presentation, antiviral mechanisms, and noncanonical NF-κB signaling (Supplementary Fig. 8C). Targeted gene set enrichment analysis (GSEA) analyses further validated the enrichment of these three processes (Figure 6C), supporting the rationale of inhibiting autophagy to improve AZA immune effects.

Given that autophagy degrades EREs, we wondered if autophagy could prevent AML blasts from being recognized by CTLs. Therefore, we segregated the 437 Leucegene AML patients based on two parameters. The first parameter was the expression of CD8A and CD8B transcripts, a reliable marker of CTL abundance in RNA-seq datasets (49). The second was the count of highly expressed AZA-induced EREs (HE-EREs), i.e., the number of EREs whose expression is above their median RNA expression across all patients having a non-null expression of the given ERE (a metric aimed at reflecting the density and diversity of epitopes possibly presented by leukemic blasts (28)) (Figure 6D). A GSEA comparing CD8^high^ vs. CD8^low^ patients within ERE^high^ and ERE^low^ groups revealed that the presence of CTLs was associated with the same processes in ERE^high^ and ERE^low^ patients, except for two gene sets related to DNA repair/proliferation, three related to metabolism, and two related to unfolded protein response and mTORC1 signaling (Figure 6E). As mTORC1 regulates autophagy, we performed additional GSEAs with four gene sets related to autophagy from the REACTOME database. We found that all four were inversely associated with the presence of CD8 T cells in ERE^high^ patients (two significantly), while the opposite was found for ERE^low^ patients (Figure 6F). Altogether, these results further support that autophagy prevents the generation of ERE-derived MAPs, thereby precluding AML blasts from being recognized by CD8^+^ T cells.

## 3. DISCUSSION

Due to their dual capacity to trigger innate immune responses and generate highly immunogenic MAPs (30), EREs represent attractive targets for developing new immunotherapeutic avenues (25,50,51). Although AZA has been proposed to promote anti-tumor CTL responses through the induction of CTA MAPs, the contribution of ERE MAPs to such responses remains elusive. Aiming to unravel this contribution, we performed a thorough proteogenomic investigation to uncover changes in the MAP repertoire after AZA treatment. As expected, we identified CTAs upregulated at the transcriptomic and immunopeptidomic levels. In contrast, ERE MAP abundance remained unchanged after AZA treatment in the four cell lines examined, suggesting that T-cell-mediated responses post-AZA treatment are more likely due to the recognition of CTA-derived MAPs than EREs. A recent report analyzing the CD8^+^ T-cell subsets targeting ERE-derived MAPs revealed no increase in ERE reactive T cells post-AZA treatment in myeloid hematological malignancies, further supporting our observations (52).

The virtual absence of ERE MAP induction by AZA was paradoxical. Indeed, AZA strongly induced ERE transcripts (Figure 2F), and the processing of numerous EREs should generate MAPs (30). In AML patients, the basal ERE expression was positively associated with the expression of molecules involved in dsRNA detection and anti-viral immune responses. In a previous report, high ERE expression in primary AML cells was associated with a favorable prognosis (53). In addition, Ohtani *et al.* demonstrated that clinical responses to AZA in myelodysplastic syndrome and AML were associated with the expression of a specific class of EREs inducing innate immune responses (54). Therefore, elevated ERE expression certainly exerts a beneficial effect on patients’ outcomes by inducing anti-dsRNA immune responses, and maximizing these responses should be pursued.

Notably, we observed that ERE levels (and associated innate immune responses) were inversely correlated to the expression of autophagy molecules in AML patients, and enhanced autophagy was found in our AZA-treated cells. Upon examination of the immunopeptidomic changes, we could attribute this latter observation to AZA’s inhibition of DNMT2 activity. Indeed, previous studies have shown that tRNA methylation by DNMT2 is involved in Asp-tRNA codon fidelity, and its loss leads to the production of misfolded proteins (55). DNMT2 inhibition is a property of AZA but not DAC. Accordingly, protein aggregation and concomitant autophagy responses were observed in AZA-but not DAC-treated AML cells. Autophagy is increasingly implicated in resistance to anti-cancer therapies, including resistance to AZA (56). We propose that autophagy mitigates tumor immunogenicity by preventing ERE MAP presentation. Interestingly, an article exploring the immunopeptidomic effects of DAC in glioblastoma cell lines evidenced an induction of ERE MAPs following similar treatment conditions to ours (57). Since DAC also induces ERE transcripts but does not induce autophagy, in contrast to AZA, we hypothesize that the autophagic process triggered by DNMT2 inhibition is responsible for the absence of ERE MAP induction following AZA treatment.

While little is known about the interplay between EREs (remnants of ancient viruses) and autophagy, it is well-reported that autophagy is a defense response against viruses. Following infection, autophagy is triggered by the signaling of pattern-recognition receptors (such as Toll-like and RIG-like receptors) (58) to sustain the presentation of intracellular source proteins by MHC-II molecules (59). Thereby, autophagy inhibition reduces the presentation of viral MAPs by infected cells to CD4^+^ T cells (60,61). In contrast, autophagy inhibition tends to increase MHC-I expression and the capacity to induce antiviral CD8^+^ T cell responses (62,63). Furthermore, autophagy can limit RIG-I-dependent IFN production by disrupting its signaling cascade (64,65). While these studies point to a potential negative correlation between autophagy and MHC-I presentation, they do not provide a precise molecular mechanism explaining how AZA-induced autophagy could prevent the generation of ERE MAPs. While this question will need to be explored in future studies, we surmise that autophagy degrades ERE RNAs instead of degrading their translational product. Indeed, autophagy has been reported to target viral (66) and ERE (34) dsRNA to autophagosomes. Notably, this would explain why CTA MAPs (which do not result from dsRNA) were successfully presented at higher levels after AZA treatment.

In conclusion, our results demonstrate that AZA-induced autophagy mitigates the ERE-dependent immune effects of AZA. They suggest that autophagy inhibition could be a desirable therapeutic option to combine with AZA. Adding autophagy inhibitors to AZA could have three desirable consequences: (1) to increase the direct cytotoxicity of AZA by preventing AML adaptation to proteotoxic stress (in agreement with results from (67)), (2) to increase ERE transcripts abundance and the subsequent beneficial anti-dsRNA innate immune responses and (3) to improve ERE MAP presentation and thereby adaptive T-cell responses. Further investigations will nevertheless be needed to verify this last hypothesis.

## ACKNOWLEDGEMENTS

The authors wish to thank Anca Apavaloaei, Jean-David Larouche, and Maria Virginia Ruiz Cuevas for valuable discussions and suggestions. We also want to thank Caroline Côté, Jalila Chagraoui, and Eugenie Goupil for technical assistance and advice regarding autophagy experiments and immunofluorescence analyses and Jeremy Zumer for advice regarding normalization methods of the mass spectrometry data. We are grateful to the IRIC genomics core facility, flow cytometry platform, Christian Charbonneau from the Microscopy facility, and Patrick Gendron from the IRIC bioinformatics platform. This study was supported by grants from the Canadian Institutes of Health Research (FDN 148400) and the Canadian Cancer Society (#705604). N.N. is supported by doctoral studentships from the IRIC and the Fonds de Recherche du Québec – Santé. J.H. is supported by post-doctoral fellowships from the Cole Foundation and the Power Corporation of Canada. BES and JC are research fellows of the FNRS. G.E. is supported by post-doctoral fellowships from the IRIC, FRQS, The Cole Foundation, and the FNRS. FB senior research associate at the FNRS Belgium.

## AUTHOR CONTRIBUTIONS

N.N., C.P. and G.E. designed the study. N.N. and G.E. performed all cell culture experiments, bioinformatic analyses, and data interpretation. G.E. built databases for mass spectrometry analysis and performed main bioinformatics analysis on differentially expressed MAPs and RNA-seq from the AML Leucegene cohort. C.D. and J.L. performed immunoprecipitation and mass spectrometry experiments. J.C. and B.E.S contributed to cell culture experiments and performed functional assays. J.H. and A.S. provided bioinformatics assistance in analyzing immunofluorescence data. M.-P.H., K.V., J-P.L., F.B., P.T., and C.P., contributed to analyzing and interpreting the data. N.N., C.P. and G.E. wrote the manuscript and the final manuscript was edited and approved by all authors.

