## Supplementary Fig. for "Autophagy degrades immunogenic endogenous retroelements induced by 5-azacytidine in acute myeloid leukemia"

### Supplementary Figures

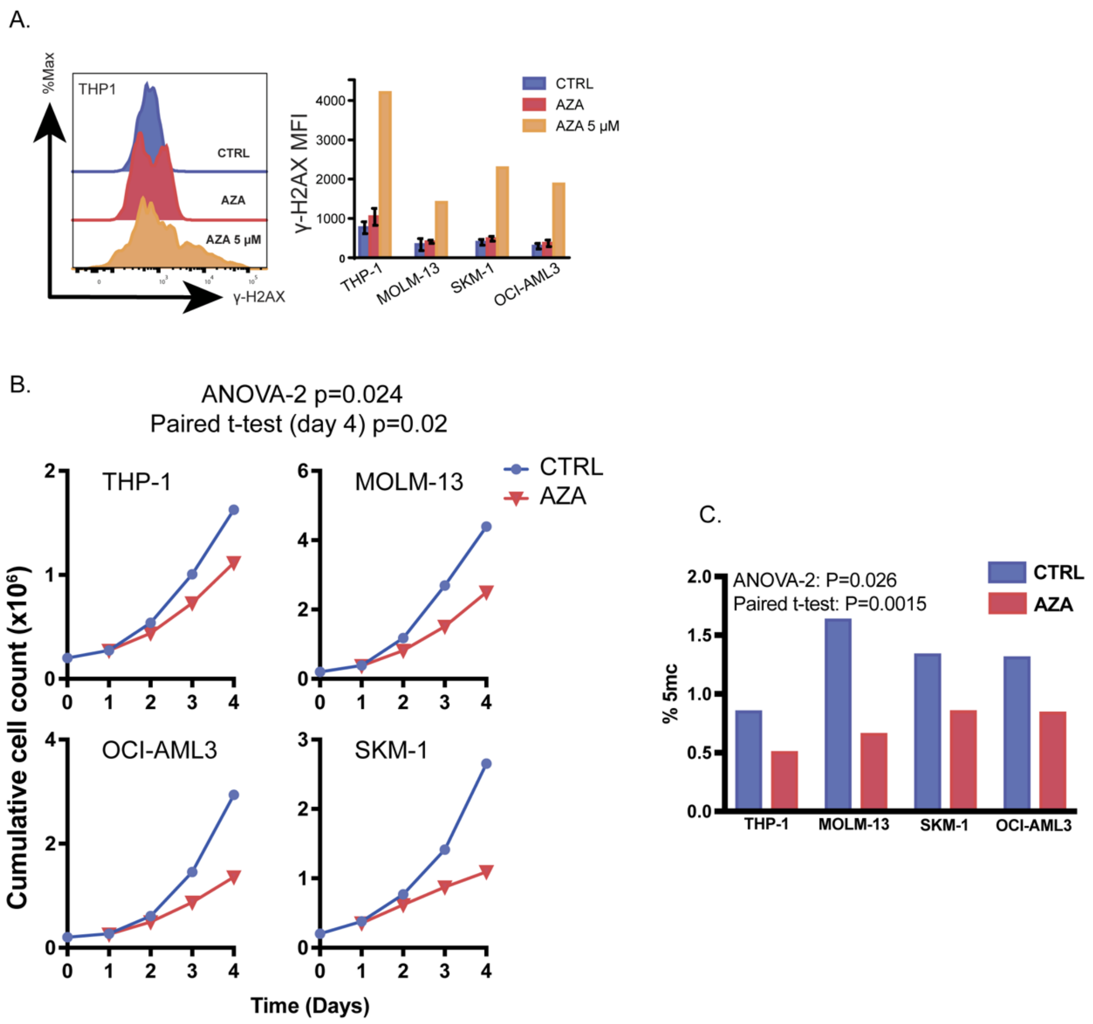


**Figure S1. Low AZA treatment reduces DNA methylation and cell growth without inducing major cytotoxic effects from DNA damage in AML cell lines**

**A**, Low AZA doses were added to four AML cell lines daily for three days (0.25 µM: MOLM-13 and SKM-1; 0.5 µM: THP-1 and OCI-AML3), and the formation of DNA double-strand breaks was monitored by flow cytometry by measuring histone H2AX phosphorylation. The left panels depict representative histograms of THP-1 cells, while the right panels depict bar plots summarizing the percentage of expression of all four AML cells. Percentages were calculated by comparing AZA-treated cells to control cells. A high AZA dose (5 µM) was used as a positive control for double-strand break formation.

**B**, Cell growth of four AML cell lines was monitored after AZA treatment by counting 7-AAD negative cells via flow cytometry

**C**, 5-methylcytosine levels measured by ELISA with the MethylFlash Global DNA Methylation Kit after AZA treatment in AML cell lines.

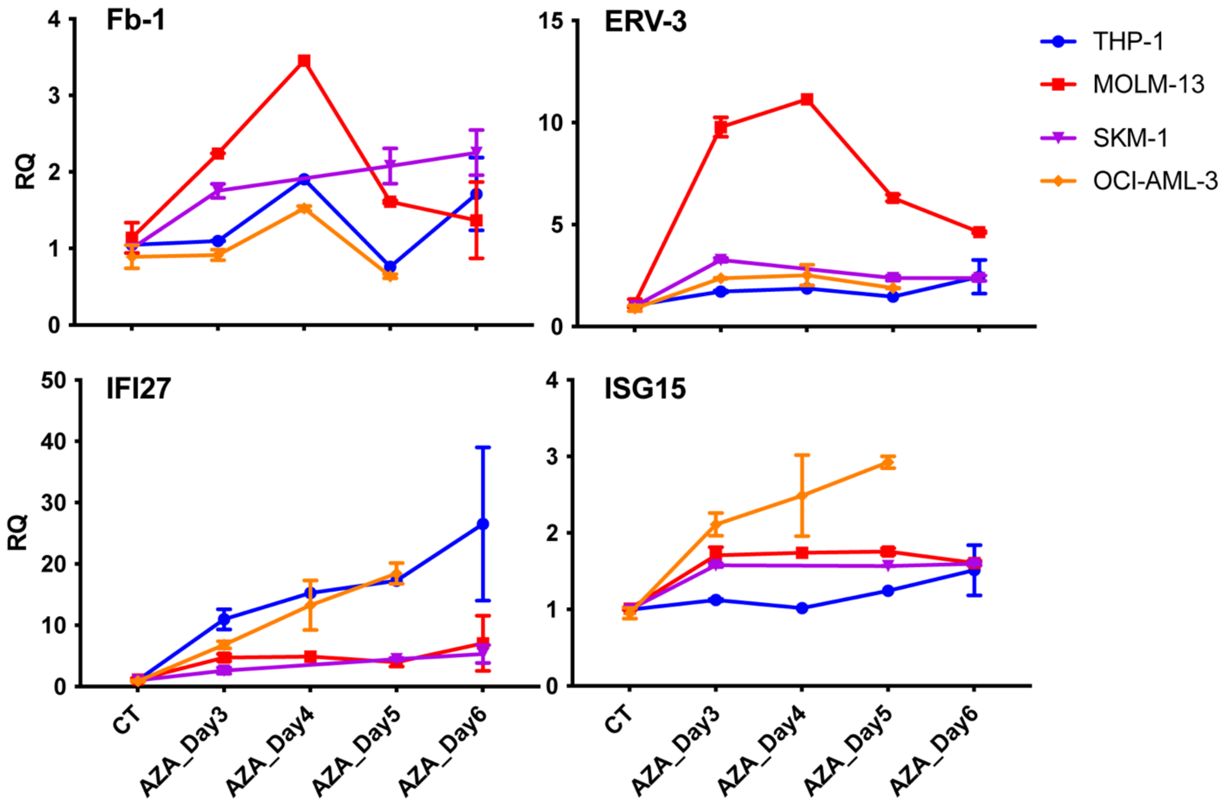


**Figure S2. Low AZA treatment leads to delayed, transient ERE and dsRNA-induced pro-inflammatory gene expression in AML cell lines**

Relative quantification levels of ERE (upper panel) and dsRNA-induced interferon (lower panel) gene candidates in four AML cell lines monitored by qPCR after AZA treatment for three days, followed by AZA discontinuation.

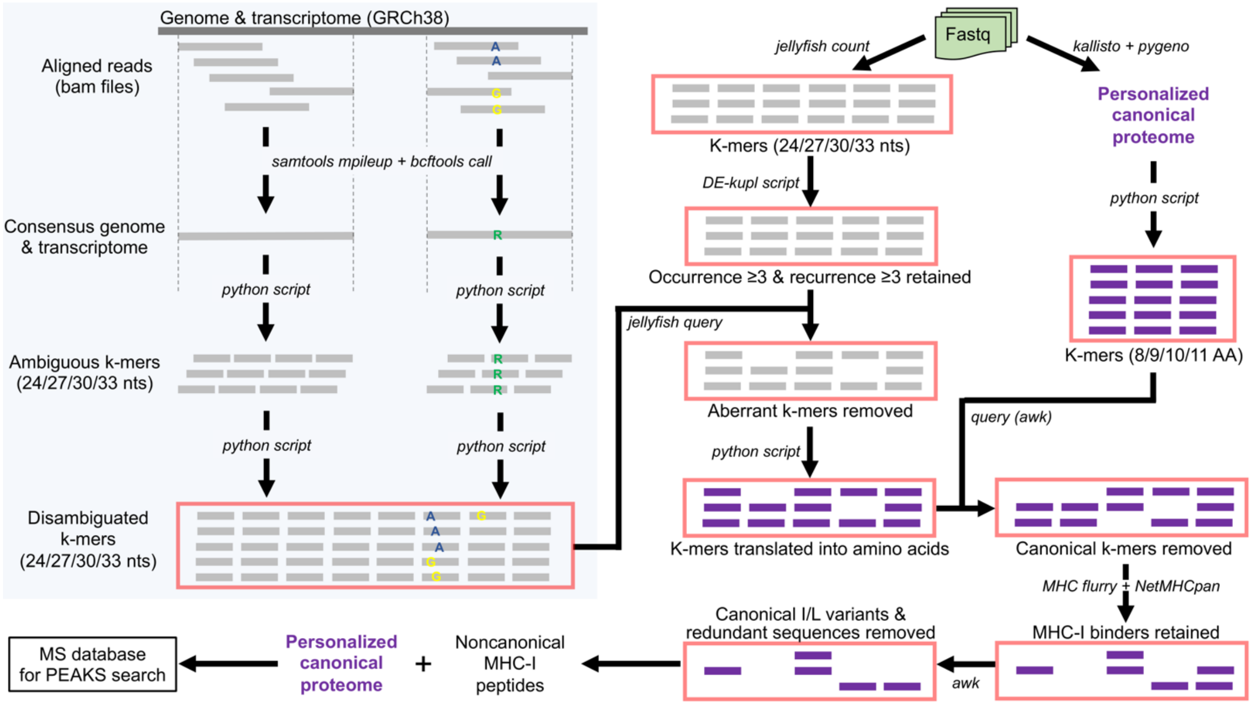


**Figure S3. Detailed proteogenomic pipeline used for database generation for MS analyses.**

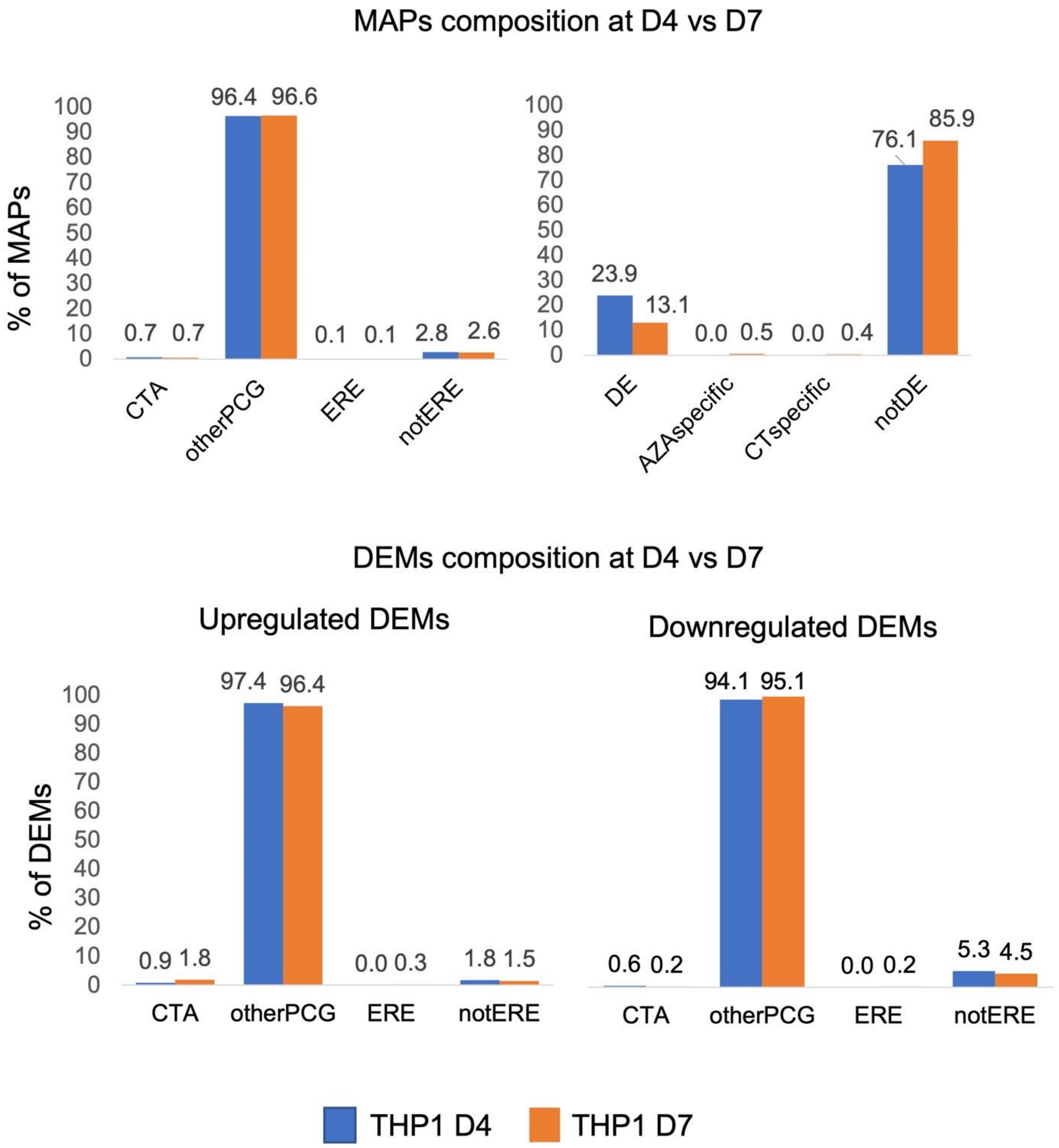


**Figure S4. Immunopeptidomic analyses at a later time-point reveal no increase in ERE-derived DEMs**

Comparison of MAP (upper panels) and DEM (lower panels) composition on days 4 and 7. OtherPCG: other protein coding genes, DE: differentially expressed.

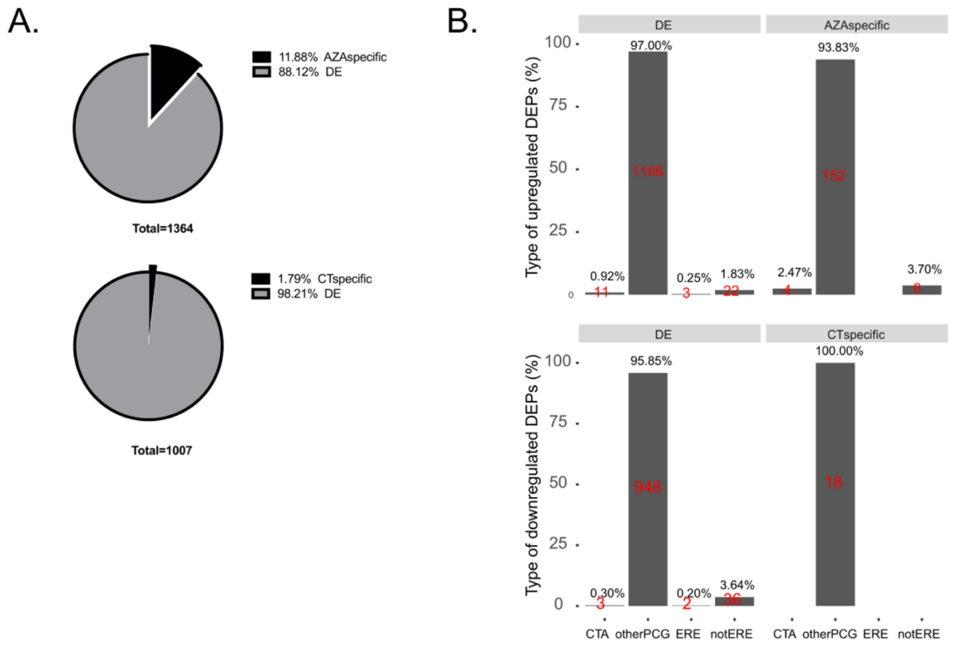


**Figure S5. MAPs presented *de novo* after treatment derived from CTAs rather than EREs**
**A**, Pie charts indicating the proportion of new MAPs previously unidentified on untreated cells (AZA specific) or MAPs unidentified after AZA treatment (CT specific) and differentially expressed for up- (upper panel) and downregulated DEMs (lower panel). **B**, Bar plots indicating DEM composition according to biotypes for up- (upper panel) and downregulated DEMs (lower panel). OtherPCG: other protein coding genes, DE: differentially expressed.

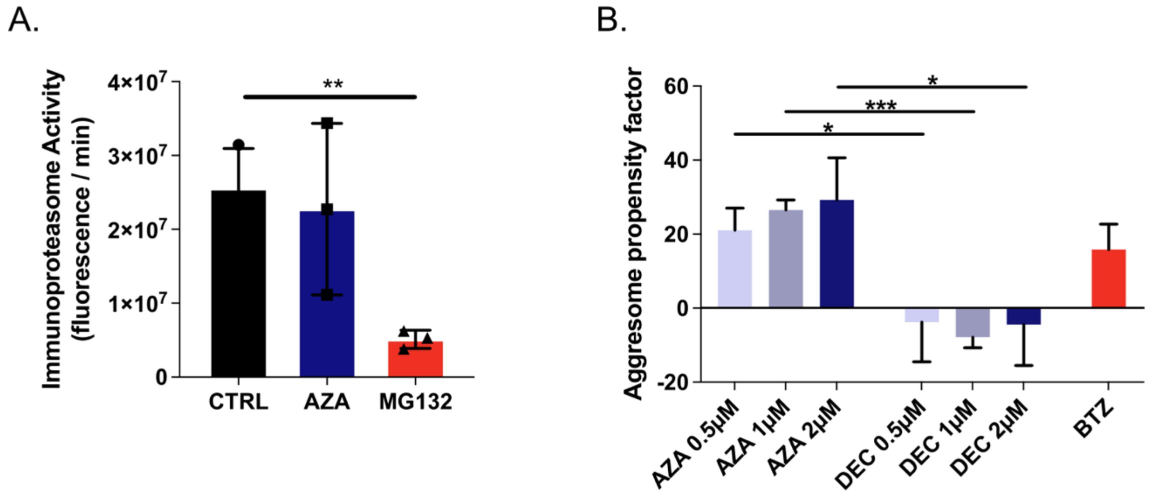


**Figure S6. AZA molds the immunopeptidome through DNMT2 inhibition**

**A**, Immunoproteasome activity monitored in THP-1 cells after AZA treatment. MG132, a proteasome inhibitor, was used as a negative control (unpaired t-test; ** p< 0. 01) **B**, Quantification of protein aggregates induced with increasing AZA and DAC concentrations in THP-1 cells. Bortezomib (BTZ) was used as a positive control (unpaired t-test; *** p< 0.001, * p< 0.05).

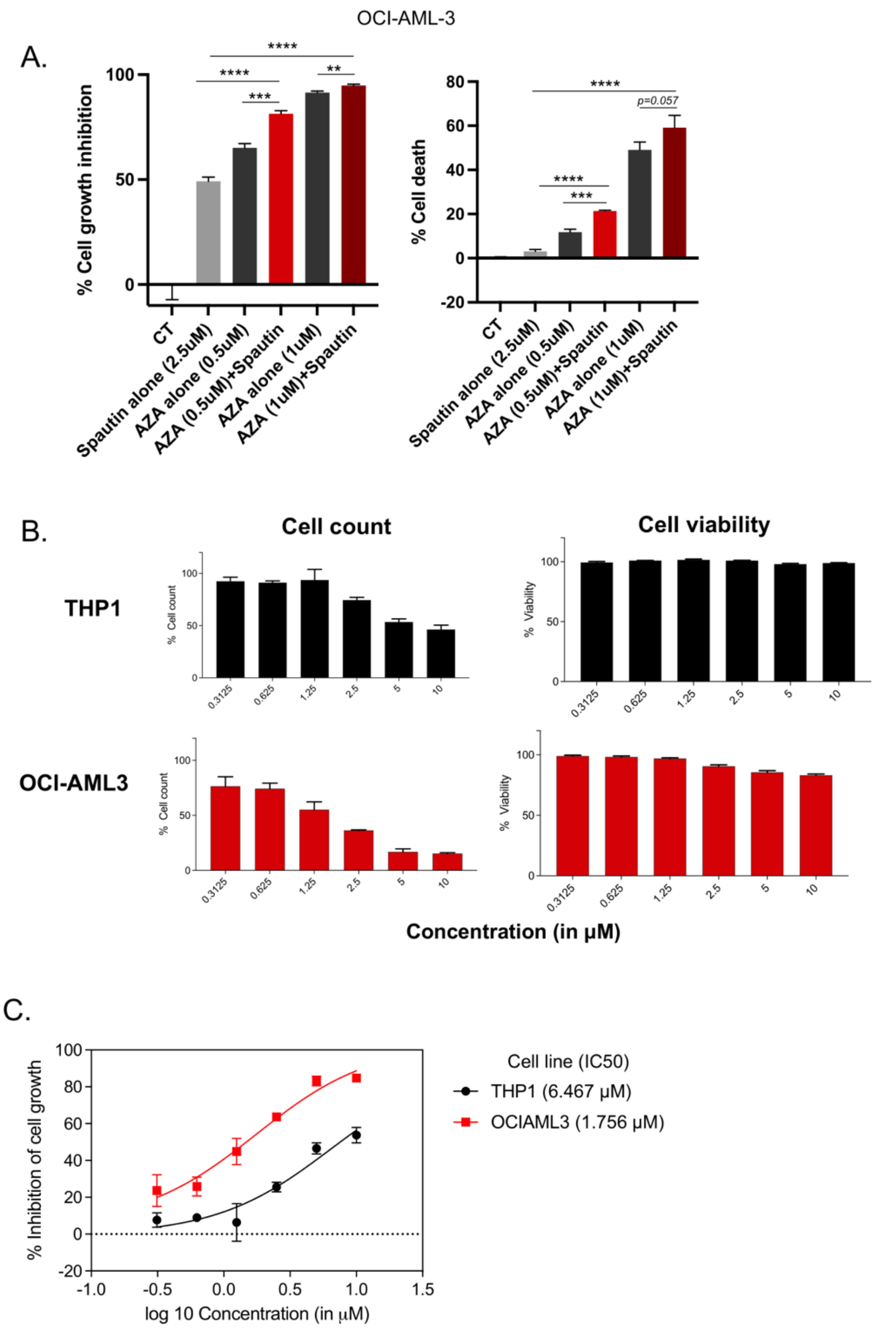


**Figure S7. Autophagy inhibition synergizes with AZA, and spautin-1 treatment alone does not induce cell death**

**A**, Cell growth inhibition and cell death of OCI-AML3 cells treated with either increasing concentrations of AZA, spautin-1, or both, monitored with 7-AAD via flow cytometry. Control cells were OCI-AML3 cells treated with 0.1% DMSO (two independent experiments; unpaired t-test; **** p< 0.0001, *** p< 0.001, ** p< 0.01). **B**, Viable cell counts after treatment with increasing concentrations of spautin-1 in THP-1 and OCI-AML3 compared to DMSO-treated control cells using 7-AAD via flow cytometry. **C**, Dose-response curves and IC50 values generated from B.

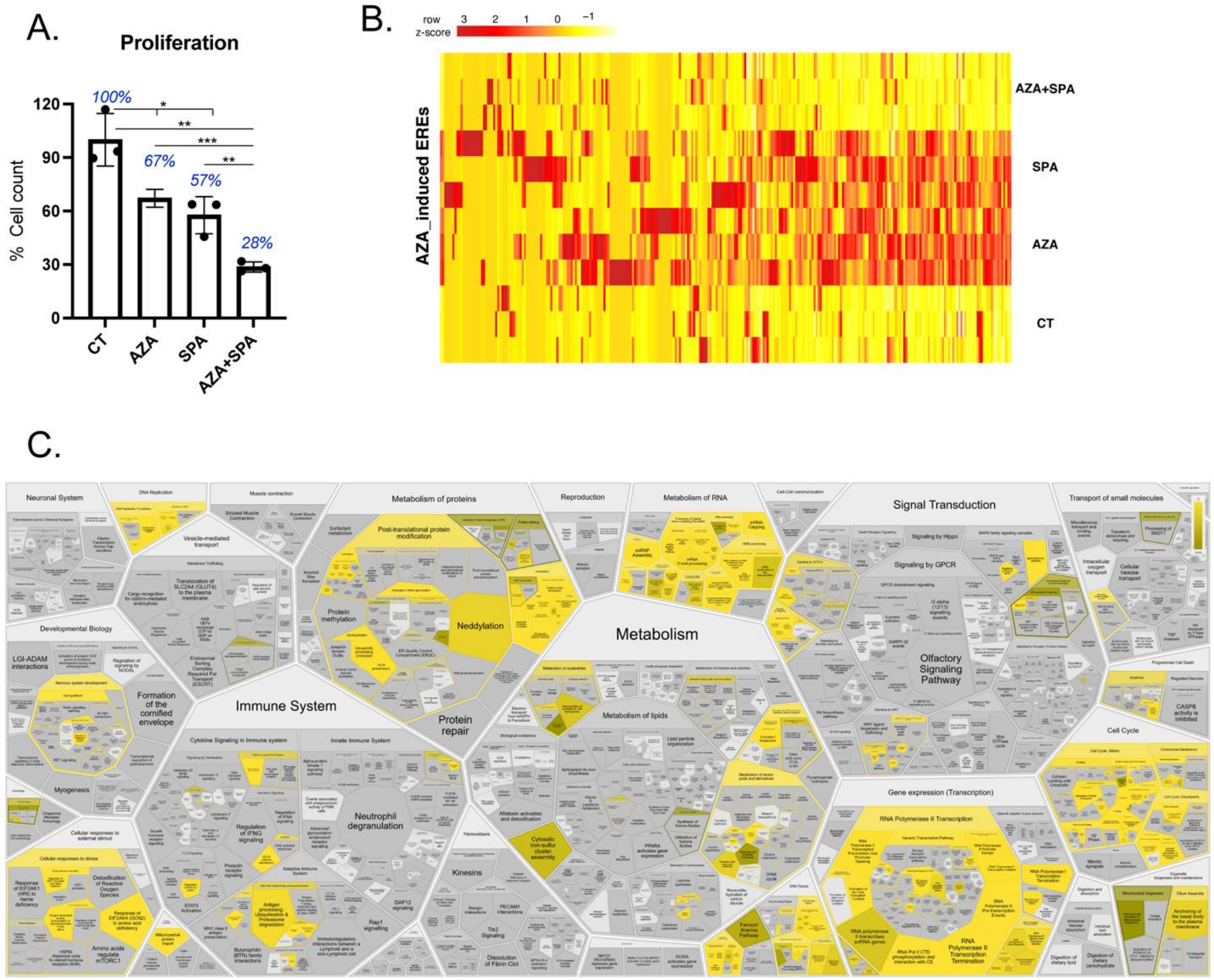


**Figure S8. Autophagy inhibition synergizes with AZA and might increase AML immunogenicity**

**A**, Cell growth inhibition with either AZA, spautin-1, or both, monitored with 7-AAD via flow cytometry in THP-1 cells (unpaired t-test; ** p< 0.01, *p< 0.05) **B**, Heatmap of AZA-induced ERE expression (in cpm) in AZA, spautin-1, AZA+spautin-1, and DMSO-treated THP-1 cells. **C**, Pathway enrichment analysis using Cytoscape software to identify biological pathways significantly enriched in AZA- and spautin-1-treated cells compared to cells treated with AZA alone. Significant pathways are highlighted in yellow.

### MATERIAL AND METHODS

#### Cell culture

THP-1, OCI-AML3, MOLM-13, and SKM-1 cell lines were freshly purchased from the Deutsche Sammlung von Mikroorganismen und Zellkulturen (DSMZ) for the current study. THP-1, MOLM-13, and SKM-1 cells were maintained in RPMI 1640 (Gibco, NY-US, 11875-093) containing L-glutamine and supplemented with 10% heat-inactivated fetal bovine serum (FBS, Gibco 12483) and 1% penicillin-streptomycin (10,000 U/mL, Gibco 15140-122). OCI-AML3 cells were maintained in MEM alpha (Gibco, NY-US, 12571063) containing L-glutamine and nucleotides supplemented with 10% heat-inactivated fetal bovine serum (FBS, Gibco 12483) and 1% penicillin-streptomycin (10,000 U/mL, Gibco 15140-122). MEM alpha without nucleotides was used to treat OCI-AML3 cells with AZA

#### Cell line treatments

For AZA treatments, cell lines were treated daily with 0.25 μM or 0.5 μM of AZA (Sigma Aldrich A2385) for 72h, followed by removal of the drug (replacement of the medium) at time points as indicated in the results section. For spautin-1 dose responses, cell lines were treated with spautin-1 (Millipore Sigma SML0440) or 0.1% DMSO control for four days, and spautin-1 or DMSO was replenished when fresh media was added. For co-treatment experiments with AZA and spautin-1 or rapamycin (kind gift from Guy Sauvageau's lab), cell lines were treated for four days with either spautin-1 (5 μM), rapamycin (0.5 μM), or 0.1% DMSO in presence of 0.5 μM of AZA. AZA was added daily for 72h, followed by discontinuation for 24h. Levels of genomic 5-methylcytosine after AZA treatment were measured by ELISA with the MethylFlash Global DNA Methylation Kit (Epigentex, P-1030).

#### Flow cytometry

Cells (~$1 \times 10^5$ cells/sample) were collected and washed 1X with PBS (Sigma P3813) before fixation/permeabilization with either the FOXP3/Transcription Factor Staining Buffer Set (eBioscience) and staining with anti-DNMT1-PE (EPR3522, Abcam) or the Fix Buffer I (Becton Dickinson, BD) followed by Phosflow Perm Buffer III (BD) and staining with anti-H2γX-AF647 (pS139, BD). All staining steps were performed at 4ºC for 30 min in the dark, and cells were pre-incubated with Fc receptor blocking antibody (BD Pharmingen 552930) for 10 min before incubation with antibodies of interest. Data were acquired on a FACS Canto II (BD).

For protein aggregate detection, $1 \times 10^5$ cells/sample were washed 3X with PBS and then fixed/permeabilized with the Cytofix/Cytoperm kit (BD) according to the manufacturer's instructions. Cells were then washed 3X with PBS and resuspended in 250 μL of assay buffer (ENZO #51035) supplemented with Proteostat Aggresome detection dye (ENZO #51035) diluted 1:10,000. Cells were analyzed with a FACS Canto II (BD) after 30 min of staining without additional washes.

Autophagy activity was measured using an autophagy assay kit (Abcam, ab139484) according to the manufacturer's protocol. In brief, cells (~$2\times10^5$/sample) were cultured for 24h in various concentrations of AZA, DAC, or rapamycin in the presence of 120 μM of chloroquine (to accumulate autophagic granules and enable their detection). Cells were collected by centrifugation and washed with assay buffer before being resuspended in 250 μL of culture medium containing 5% FBS and mixed with 250 μL of diluted green staining solution. Cells were incubated for 30 min at 37ºC in the dark and washed with assay buffer. Relative autophagy activities were measured using a Cytoflex flow cytometer (Beckman Coulter).

**Immunoproteasome activity**

Immunoproteasome activity was assessed on THP-1 or OCI-AML3 cells (~$5\times10^5$ cells/sample) lysed in 1 mL of lysis buffer (50 mM Tris-HCl, 2 mM DTT, 5 mM $MgCl_2$, 10% (v/v) glycerol, 2 mM ATP, and 0.05% (v/v) digitonin). The assay was performed immediately after lysis with the Immunoproteasome Activity Fluorometric Assay Kit I (Ubiquitin-Proteasome Biotechnologies, TX-US, J4160) according to the manufacturer's instructions. Fluorescence was detected using the TriStar² LB 942 microplate reader (Berthold Technologies GmbH & Co.KG).

**Real-time PCR**

Quantitative real-time PCR was performed for candidate ERE and dsRNA-induced interferon genes using validated Universal ProbeLibrary assays (Roche) on the Viia7 Real-time PCR system (Applied Biosystems). Relative target mRNA levels were normalized to GAPDH and ACTIN and analyzed using Expression Suite software v1.1 (ThermoFisher).

**Immunofluorescence experiments**

THP-1 cells were attached on chambered slides (iBidi 80826) with poly-L-lysine and fixed using ice-cold methanol for 15 min at -20ºC, washed three times with PBS, and incubated with saturation buffer (5% BSA-PBS) for 1h. Primary antibody was added (1:200, anti-dsRNA, clone J2, SCICONS) and incubated overnight at 4ºC. Cells were washed three times for 15 minutes with PBS on a shaker, followed by incubation with secondary antibodies (1:2000 goat anti-mouse IgG AlexaFluor 594 Invitrogen A-11020) at room temperature for 1h and washed three times for 10 minutes with PBS. Next, cells were incubated with DAPI containing PBS, and slides were stored at 4ºC in the dark for at least three days before confocal analyses. Confocal analyses were performed with a Zeiss LSM700 confocal microscope, and images were quantified using EBImage package on R. Transfected cell lines with Poly(I:C) were used as positive controls. No unspecific staining was observed with secondary antibodies alone.

**Library preparation and RNA sequencing**

Total RNA was isolated using TRIzol (Thermo Scientific) followed by RNeasy purification (Qiagen). RNA was quantified using Qubit (Thermo Scientific), and quality was assessed with the 2100 Bioanalyzer (Agilent Technologies). Transcriptome libraries were generated using the KAPA RNA HyperPrep (Roche) using a poly-A selection (Thermo Scientific). Sequencing was performed on the Illumina NextSeq500, obtaining around 120M paired-end reads per sample (60M clusters) for AZA vs. control in AML cell lines and 30M single-end reads per sample for spautin-1+AZA validation RNA-seq experiment. Raw data were deposited on GEO (GSE217572).

**Gene expression analyses**

All transcript expression (canonical genes and EREs) quantifications were performed with kallisto v0.43.0 (1) with default parameters. Kallisto's transcript-level count estimates were converted into gene-level counts using the R package tximport. EdgeR was used to normalize counts using the TMM algorithm and output count-per-million (cpm) values. Differential gene expression analyses were conducted in R3.6.1, as reported previously (2). In brief, raw read counts were converted to cpm, normalized relative to library size, and lowly expressed genes were filtered by keeping genes with cpm >1 in at least two samples using edgeR 3.26.8 (3) and limma 3.40.6 (4). Subsequently, voom transformations and linear modeling using limma's lmfit were performed. Moderated t-statistics were then computed with eBayes. Genes with false-discovery rates ≤0.05 and $-1 \geq \log_2(FC) \geq 1$ were considered significantly differentially expressed. For differential gene expression analyses performed on AZA-treated cell lines, a unique paired analysis comparing AZA-treated vs. control cells was performed.

Gene ontology and biological pathway annotations were performed with DAVID v6.8 (https://david.ncifcrf.gov). Functional annotations with a p-value <0.05 were considered significant. Gene set enrichment analysis (GSEA) was performed with the fgsea package in R (5). A pre-ranked gene list was generated by ranking expressed genes obtained from limma-voom on moderated t-statistics. Gene sets were obtained either from the HALLMARK or REACTOME matrix, downloaded from the MSigDB database. The enrichment analysis for REACTOME gene sets among genes significantly upregulated in the AZA+Spautin-1 vs. AZA-only cells was performed with the Reactome FI module in Cytoscape v3.7.2 (6).

**Database generation for mass spectrometry identifications**

To build databases to analyse MAPs originating from any region of the genome (canonical exons, introns, EREs, ncRNAs, intergenic regions, etc.) and including MAPs deriving from mutations present in the genome of the analyzed cell line, we adopted an alignment-free proteogenomic approach. We built two personalized, non-overlapping proteomes, canonical and non-canonical, and concatenated them to perform MS identifications. All scripts and usage instructions for the pipeline can be found on Zenodo (doi 10.5281/zenodo.7096388).

*Personalized canonical proteomes*

RNA-Seq reads were trimmed using Trimmomatic v0.35 and aligned to GRCh38.88 using STAR v2.5.1b (7) running with default parameters except for --alignSJoverhangMin, --alignMatesGapMax, --alignIntronMax, --quantMode and --alignSJstitchMismatchNmax parameters for which default values were replaced by 10, 200,000, 200,000, TranscriptomeSAM and "5 -1 5 5", respectively, to generate bam files. Single-base mutations with a minimum alternate count setting of 5 were identified using freeBayes 1.0.2-16-gd466dde (8). Transcript expression was quantified in transcripts per million (tpm) with kallisto v0.43.0 with default parameters. Finally, we used pyGeno (9) to insert high-quality sample-specific single-base mutations (freeBayes quality >20) into the reference exome and export sample-specific sequences of known proteins generated by expressed transcripts (tpm >0) to generate fasta files of personalized canonical proteomes.

*Personalized non-canonical proteomes*

Step 1. We built consensus genomes and transcriptomes (including only genomic regions covered by RNA-seq reads and single-nucleotide polymorphisms as ambiguous nucleotides) from STAR-generated bam files of each sample (per replicate per cell line). This was performed with the reference genome and transcriptome as input of the samtools (10) and bcftools suites (11): 'samtools mpileup -C50 -uf reference.fasta sample.bam | bcftools call -c | vcfutils.pl vcf2fq | gzip >> consensus.fastq.gz'. The consensus genomes and transcriptomes were then chopped into k-mers (of 24, 27, 30, or 33 nucleotide lengths, corresponding to the length of MAPs: 8–11 amino acids) with a homemade python script, and k-mers containing consensus nucleotides (R|Y|M|K|W|S|B|D|H|V|N) were disambiguated (A|T|C|G) using a homemade python script. These k-mers were then reverse-complemented, and all k-mers (non-ambiguous and disambiguated, originals and reverse-complemented) were assembled in a single database generated with Jellyfish v2.2.3 (12).

Step 2. The fastq files of each sample (per replicate per cell line) were used to generate k-mer libraries of either 24, 27, 30, or 33 nucleotides in length containing k-mers present at least twice per sample. This was performed with Jellyfish: 'jellyfish count -L 2 -m <length> -F 2 -s 1G -o sample.jf' on trimmed forward and reverse-complemented (with the fastx_reverse_complement function of the FASTX-Toolkit v0.0.14) reverse fastq files. Next, the k-mer databases were combined into single databases per cell line (four databases were obtained eventually, one per MAP length) by keeping only those k-mers with three occurrences in three different samples (out of six samples: three controls and three AZA-treated). This was performed with a script ('joinCounts') obtained from the DE-kupl pipeline (13): 'joinCounts -r 3 -a 3 <fastq files>'. This allowed us to retain k-mers that most likely generate MAPs (since high RNA expression is a robust predictor of MAP generation (14,15)).

Step 3. The k-mers generated in step 2 were queried in the k-mer databases generated in step 1. This allowed us to discard consensus artifacts such as exon-intron junctions, wrong SNP calling, false intron coverage and to filter k-mers on their minimum occurrence and inter-sample sharing. The query was performed with the 'jellyfish query -i' command.

Step 4. The personalized canonical proteomes were chopped into peptide k-mers (8, 9, 10, or 11 amino acids) using a homemade python script.

Step 5. The resulting k-mers from step 3 were translated into their peptide sequence with a homemade python script. Peptide k-mers containing stops were removed (with awk), and peptide k-mers generated in step 4 were removed from this list to prevent overlaps between the canonical and non-canonical proteome.

Step 6. The non-canonical peptides were tested for their capacity to bind HLA alleles of their respective cell line (determined with Optitype (16)) with either MHC flurry 1.4.0 (17) or netMHCpan 4.0 (18) for alleles not supported by MHC flurry. Predictions were made with the epitopepredict module (19) to handle MHC flurry and NetMHCpan. Peptides with a percentile rank ≤2% were kept for further processing.

Step 7. Since leucine and isoleucine variants are not distinguishable by standard MS approaches, we inspected the list of non-canonical peptides and discarded those for which an existing variant (MHC binder as well) was flagged as canonical. Next, short peptides with sequences completely included in the sequence of longer peptides were discarded (awk) from the list, and peptide sequences were used to generate a fasta file, eventually concatenated with the personalized canonical proteome to generate the final MS databases.

**MHC-I peptide isolation by immunoprecipitation**

W6/32 antibodies (BioXcell) were incubated in PBS for 60 min at room temperature with PureProteome protein A magnetic beads (Millipore) at a ratio of 1 mg of antibody per 1 mL of slurry. Antibodies were covalently cross-linked to magnetic beads using dimethylpimelidate as described (20). The beads were stored at 4ºC in PBS (pH 7.2) and 0.02% $NaN_3$. Frozen cell pellets (118–135×$10^6$ cells/pellet) were thawed and resuspended in 0.4 mL PBS (pH 7.2) and solubilized with 1 mL of detergent buffer containing PBS (pH 7.2) and 1% (w/v) CHAPS (Sigma) supplemented with a protease inhibitor cocktail (Sigma). Cell pellets were incubated for 60 min with tumbling at 4ºC and then spun at 16,600g for 20 min at 4ºC. Supernatants were transferred into new tubes containing 1 mg of W6/32 antibody covalently-cross-linked protein A magnetic beads per sample and incubated with tumbling for 20h at 4ºC. Samples were placed on a magnet to recover bound MHC-I complexes to magnetic beads. Magnetic beads were first washed with 8× 1 mL PBS, then with 1× 1 mL of 0.1X PBS, and finally with 1× 1 mL of $H_2O$. MHC-I complexes were eluted from the magnetic beads by acidic treatment using 0.2% formic acid (FA). To remove residual magnetic beads, eluates were transferred into 2.0 mL Costar mL Spin-X centrifuge tube filters (0.45 mm, Corning) and spun for 5 minutes at 855g. Filtrates containing peptides were separated from MHC-I subunits (HLA molecules and β-2 macroglobulin) using homemade stage tips packed with two 1 mm diameter octadecyl (C-18) solid phase extraction disks (EMPORE). Stage tips were pre-washed with methanol, then with 80% acetonitrile (ACN) in 0.1% trifluoroacetic acid (TFA), followed by 0.1% TFA, and finally with 1% TFA. Samples were loaded onto stage tips and washed with 1% TFA, followed by 0.1% TFA. Peptides were eluted with 30% ACN in 0.1% TFA, dried using vacuum centrifugation, and then stored at -20ºC until MS analysis.

**TMT labeling**

MHC-I peptide extracts were reconstituted in 200 μL of 200 mM HEPES buffer (pH 8.2). TMT0-126 reagents or TMT6-plex (Thermo Fisher Scientific) were dissolved in 40 μL of anhydrous ACN (Sigma-Aldrich), and 5 μL of 0.02 mg/ μL was added to the peptides. The solutions were gently mixed and incubated for 90 min without agitation at room temperature before the reactions were quenched by hydroxylamine (Thermo Fisher Scientific). Samples were desalted on homemade C18 stage tips and dried down.

**Mass spectrometry analyses**

Dried peptide extracts were resuspended in 4% FA (EMD Millipore) and loaded on a custom C18 analytical column (20 cm × 150 mm i.d. packed with C18 Jupiter Phenomenex) with a 106-min gradient from 0% to 30% ACN (0.2% FA) and a 600 nL/min flow rate on an EasynLC II system. Samples were analyzed with an Exploris mass spectrometer (Thermo Fisher Scientific) in positive ion mode with the source at 2.8 kV. Each full MS spectrum, acquired with 240,000 resolution, was followed by MS/MS spectra, where the most abundant multiply charged ions were selected for MS/MS sequencing with a resolution of 30,000, 100% normalized automatic gain control, injection time of 700 ms, and collisional energy of 36%.

**Identification of MAPs and differential MAP analyses**

Liquid chromatography (LC)-MS/MS (LC-MS/MS) data were searched against respective cell line-specific databases using PeaksXPro. For peptide identification, no enzyme was selected, and tolerance was set at 10 ppm and 0.01 Da for precursor and fragment ions, respectively. The occurrences of oxidation (M) and deamidation (NQ) were set as variable modifications. Following peptide identification, we used a modified target decoy approach built-in PEAKS and applied a sample-specific threshold on the PEAKS score to ensure a false discovery rate of 5%, calculated as the ratio between the number of decoy hits and the number of target hits above the score threshold. Binding affinities to the sample's HLA alleles were predicted with NetMHCpan 4.1b (21), and only 8 to 11-amino-acid-long peptides with a rank eluted ligand threshold ≤ 2% were used for further annotation; these filtering steps were performed with MAPDP software (22). Intensities of all modifications for a single peptide were summed, and peptides containing too many missing values were eliminated by keeping peptides quantified in two out of three replicates of at least one condition. Next, VSN normalization was performed, which was the best available normalization method based on analyses with NormalyzerDE (23). Imputation for missing values was performed by Perseus with width of 0.3 and downshift of 1, and MAPs with p-values <0.05 and fold-changes (FC)>2 were considered significantly differentially expressed using limma analysis. MAPs exclusively detected in one condition were defined by having valid values from all three biological replicates in one condition while no values in the other condition.

**Biotype attribution to identified MAPs**

BamQuery (24) was used to annotate if MAPs derived from protein-coding, EREs, or other non-coding regions. CTAs were annotated using (25).

**Bioinformatic analyses performed on MAPs**

Amino acid compositions, aromaticity, and GRAVY indexes were assessed with the ProtParam module of Biopython. The RNA expression of each MAP was obtained using BamQuery (24).

**Quantification and statistical analysis**

Unless indicated otherwise, all statistical tests comparing two conditions were performed using the Mann–Whitney U test. All correlations were assessed with the Pearson correlation coefficient. Unless mentioned otherwise, all boxes in boxplots represent the median, $25^{th}$, and $75^{th}$ percentiles, and whiskers extend to the 10th and 90th percentiles. Unless mentioned otherwise, all bar plots represent the average with standard deviation (SD). Plots and statistical tests were mainly performed with GraphPad Prism v9.1.1. For all statistical tests, **** refers to $p< 0.0001$, *** refers to $p< 0.001$, ** refers to $p< 0.01$, and * refers to $p< 0.05$.

**Data and Code Availability**

In-house scripts used in this study are available on Zenodo at DOI: 10.5281/zenodo.7096388. All other sequencing and expression data have been deposited to the NCBI Sequence Read Archive and GEO under accession code #GSE217572.
